# ACCURATE PREDICTION OF ASPARAGINE DEAMIDATION IN BIOLOGICS USING ADVANCED MACHINE LEARNING MODELS

**DOI:** 10.64898/2026.01.06.697962

**Authors:** Shafayat Ahmed, Nicole Swope, Valentin Stanev, Romina Hofele, Dominique WuDunn, Rohan Jain, Jared Delmar, Maryam Pouryahya

## Abstract

The spontaneous deamidation of asparagine residues remains a major obstacle to the stability and efficacy of protein therapeutics. Currently available models in the literature for predicting deamidation liabilities can suffer from limited generalizability, likely due to biases such as sequence similarity within datasets. In this study, we built machine learning models using protein language models (e.g., ESM2) and graph neural networks (GNNs), trained on a comprehensive dataset of 591 asparagine sites from over 105 protein molecules. To address the critical issue of data leakage, we implemented a peptide grouping strategy yielding more accurate estimates of model performance for novel deamidation sites. Our analysis shows that, when sequence similarity bias is controlled, protein language models match traditional feature-based models that use amino acid composition, k-mers, PSSMs, and predicted secondary structure/solvent accessibility, while offering substantial computational advantages. Additionally, our GNN-based pipeline further increases prediction accuracy by up to 8% compared to language model-only tools and delivers a 15–25% improvement over motif-based approaches. This methodological framework enables more reliable and rapid in-silico prediction of deamidation liabilities, potentially reducing costly late-stage interventions in protein therapeutic development and is generalizable to the modeling of additional protein post-translational modifications.

## 1 Introduction

Protein therapeutics are a rapidly growing pharmaceutical segment, now accounting for about half of recent biologic drug approvals [21]. This class which includes monoclonal antibodies, antibody-drug conjugates, fusion proteins, enzymes, and vaccines offers new treatment options for previously intractable diseases, but also faces unique development challenges, notably chemical degradation risks. Spontaneous deamidation of asparagine residues, forming aspartic or iso-aspartic acid, is among the most common, impacting drug activity, pharmacokinetics, and immunogenicity [17, 1, 7]. Accurate prediction and mitigation of asparagine deamidation is therefore critical for successful drug development.

Computational tools for deamidation prediction have evolved from sequence rule-based models that identify susceptible NG, NS, and NN motifs [18] to structure-informed and machine-learning approaches that exploit predicted or homology-derived structural features [9, 22, 16, 12, 20, 3, 8, 5]. Nevertheless, current models are limited by small training datasets and methodological challenges associated with homology modeling. In addition, inadequate handling of sequence similarity between training and test sets has frequently led to inflated performance estimates. This problem is particularly acute for antibody and antibody-derived therapeutics, where conserved or similarly engineered regions are common; random cross-validation can place related sequences in both training and validation/test partitions, resulting in data leakage and overstated metrics. Many prior models have been trained on commercially available or in-house biologics that exhibit substantial within-dataset sequence similarity, further exacerbating these issues.

Recent advances–most notably AlphaFold2 for structure prediction [10] and protein language models such as ESM2 for deamidation prediction [13] offer promising avenues to mitigate these limitations. AlphaFold2 enables structural modeling in the absence of suitable templates, while protein language models leverage large-scale sequence corpora to infer functional properties without explicit structural inputs, potentially enhancing generalizability, robustness, and computational efficiency.

We developed an improved framework for deamidation prediction by curating a large unbiased dataset and preventing data leakage, showing that protein language models offer reliable, scalable assessment for protein therapeutics. To summarize our main contributions:

- **Rigorous Data Integrity Assessment:** We thoroughly investigated the influence of data splitting strategies on prediction validity, demonstrating that grouping sequence-similar peptides in the same fold is essential for preventing data leakage and yielding accurate generalization metrics. We also quantified how different sequence window sizes surrounding Asn sites affect model sensitivity and specificity, supporting rational parameter selection.
- **Comprehensive Model Pipeline Development:** We constructed and evaluated four major prediction pipelines, leveraging traditional sequence- and structure-based features to establish baseline performance and integrating state-of-the-art protein language models–such as ESM2–with graph neural networks. This framework enables learning of complex, context-dependent relationships directly from both sequence and structural attributes. Across these approaches, we systematically explored input feature sets, protein language model variants, and graph neural network configurations to optimize predictive accuracy.
- **Practical Implications:** Our work establishes a robust computational framework capable of rapid, scalable, and reliable in-silico deamidation liability assessment. This significantly reduces the risk of late-stage pipeline failure due to chemical instability and enables its integration into early-phase screening of biopharmaceutical candidates.

## 2 Experiments and Data Collection

### 2.1 Sample Preparation and LC-MS/MS Analysis

Therapeutic protein samples at 10 mg/mL were incubated at 40°C and pH 6.0 for up to 4 weeks to accelerate deamidation. This stress condition was selected as more representative of manufacturing and storage conditions compared to the pH 8.0 conditions used in our previous study [4]. Samples were collected at 0, 2, and 4-week timepoints and stored at -80°C prior to analysis.

LC-MS/MS tryptic peptide mapping was performed following the protocol described in our previous work [4], with modifications for improved site localization. Briefly, samples were denatured, reduced, alkylated, and digested with trypsin before separation on a reversed-phase C18 column and analysis by high-resolution mass spectrometry. Deamidation quantitation was based on extracted ion chromatography peak areas, with site-specific assignments confirmed by MS/MS fragmentation when possible.

### 2.2 Protein Structure Generation and Feature Extraction

Full-length protein structures were generated using AlphaFold2 Multimer (v2.3.2) [6], representing a significant advancement over the homology modeling approach used in our previous work [4, 3]. This method enables accurate structure prediction for molecules lacking suitable homologous templates in the Protein Data Bank, particularly for non-antibody proteins. For monoclonal antibodies, structures were truncated to the variable fragment (Fv) region to prevent the appearance of redundant conserved asparagine sites in our training dataset, while retaining all valuable variable region asparagine sites relevant to deamidation prediction. For the structure-based model, features were extracted for each asparagine residue using custom Python scripts utilizing mainly the Biopython package (v1.80), following a similar framework established in our previous study, but significantly augmented to include additional valuable features. These parameters included: (1) N+1 residue identity as both a categorical variable and empirical pentapeptide half-life (pphl) [19], (2) backbone dihedral angles (phi and psi) of N, N+1, and N-1 residues, (3) asparagine side chain dihedrals (chi1 and chi2), (4) nucleophilic attack distance between the side chain carbonyl and backbone nitrogen, (5) solvent accessibility expressed as both percentage and absolute area, (6) hydrogen bonding to the asparagine side chain, (7) secondary structure classification of N, N+1, and N-1 residues, (8) IMGT properties of N+1 and N-1 residues, including volume, hydrophobic moment, isoelectric point, polarity, and charge [15], and (9) local formal charge in the vicinity of the backbone N+1 residue nitrogen. These parameters captured the relevant local structural environment and neighboring residue properties as described previously [3].

### 2.3 Data Quality Control and Preprocessing

Rigorous data cleaning procedures were implemented to ensure training set quality. Asparagine sites with ambiguous mass spectrometric assignments were excluded from the dataset. This included cases where multiple asparagines occurred within a single tryptic peptide without sufficient MS/MS fragmentation data to localize the modification site. Additionally, sites from peptides with known complex deamidation patterns (such as the conserved “PENNY” motif [2]) were handled based on extensive experience with this conserved peptide in order to quantify individual site contributions from peptide-level measurements. Asparagine residues followed by proline (“NP sites” where N+1=P) were removed from the dataset, which is a preprocessing step not seen in the literature or our previous publication. This is important because NP sites cannot undergo deamidation; the absence of a backbone nitrogen on proline prevents nucleophilic attack on the Asn side chain carbonyl carbon atom and subsequent succinimide ring formation. As all NP sites will belong to the negative deamidation class, and the backbone structure is significantly different and unique from all other sites, removal of these sites improves learning of the model on relevant sites and structures [8]. Deamidation rates were calculated by least-squares fitting to an exponential decay function, and sites were classified as deamidating (TRUE) or non-deamidating (FALSE) based on a threshold of both ≥2.0% and ≥5.0% deamidation per month under stress conditions. In total, the dataset comprises 591 sites, of which 485 are negative (non-deamidating) and 106 are positive (deamidating) according to the 5.0% cutoff threshold (see Figure 1a).

**Figure 1:**
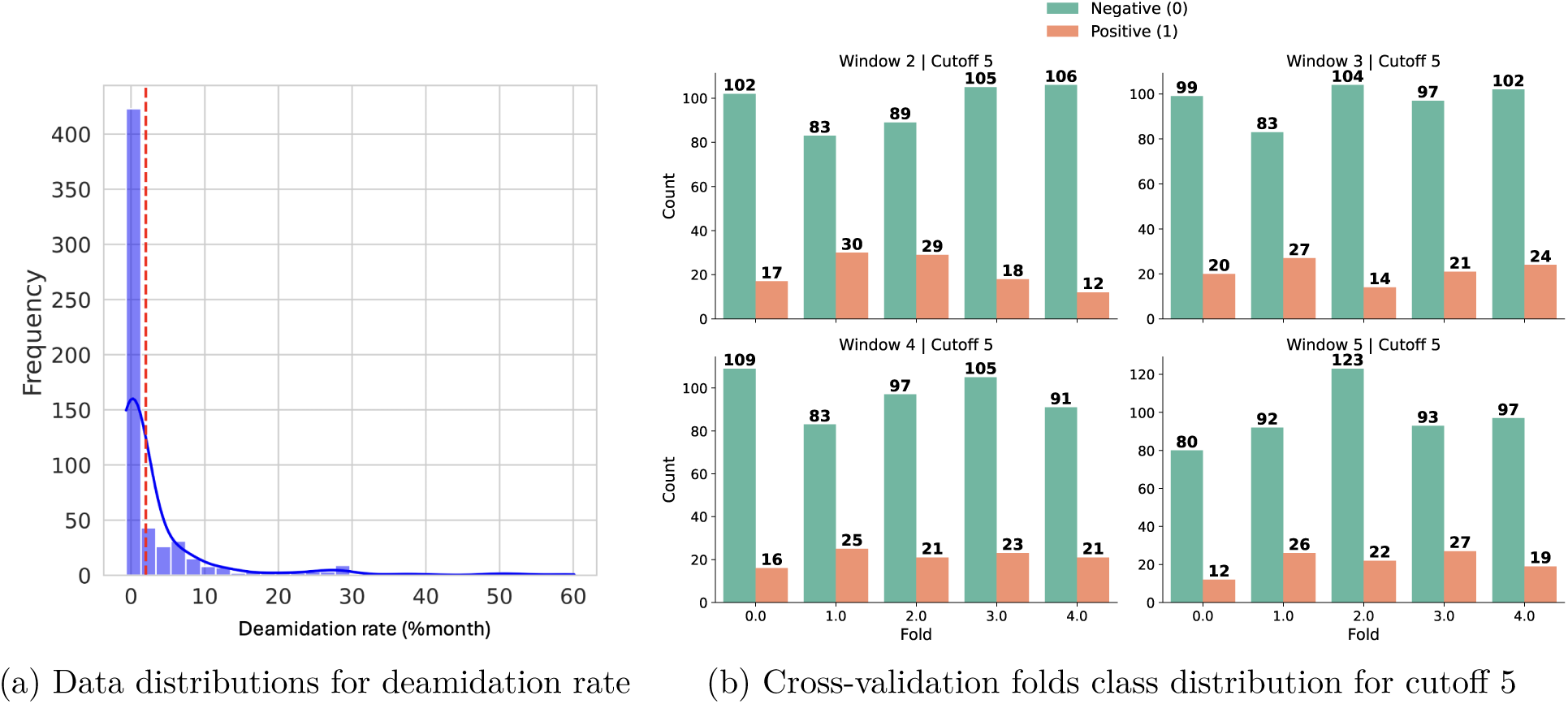
Data distribution of the collected samples and their deamidation rates. In panel (a), problematic antibodies are characterized by higher values further from zero, as indicated by the red dashed line. Panel (b) shows the count of positive and negative samples per fold for deamidation cutoff rate 5, illustrating the sample balance across windows and experimental groups.

## 3 Methodology

### 3.1 Problem Formulation

We frame deamidation prediction as a binary classification task. Each sample corresponds to an asparagine residue, labeled as positive if its experimentally measured deamidation rate exceeds the cutoff (≥2.0% and ≥5.0%) and negative otherwise.

### 3.2 Data Splitting Strategies

Careful data splitting is essential for fair evaluation of predictive models. If information from the test set leaks into training, or if highly similar samples appear in both sets, performance may be inflated. Conversely, suboptimal splitting can underestimate accuracy when splits are unbalanced or unrepresentative.

#### 3.2.1 Peptide Group Data Splitting

We defined a window size to specify the local sequence context used to group residues for the train–test split. Each window is centered on the target asparagine (N) and grows outward by alternately extending to the N-terminus and C-terminus of the sequence. For example, size 1 includes [N, N+1], size 2 expands to [N-1, N, N+1], size 3 to [N-1, N, N+1, N+2], and size 4 to [N-2, N-1, N, N+1, N+2], and so on. If multiple residues have the same sequence within their defined window, they are grouped together into clusters. Each cluster is used as the basic grouping for assigning samples to cross-validation folds, preventing highly similar local sequence contexts from appearing in both the training and test sets.

Window size directly influences the risk of information leakage between training and test sets in our clustering approach. Smaller windows, which represent short sequence motifs around the asparagine, are more likely to recur across different peptides, resulting in larger clusters that group together many residues sharing the same local context. By assigning each cluster wholly to either the training or test set, we ensure that identical local motifs do not span both splits and thus strictly minimize data leakage. Conversely, larger windows capture extended and specific sequence contexts, leading to smaller clusters with less recurrence. In this scenario, key local motifs may be distributed across different clusters and assigned to both training and test sets, increasing the possibility for information leakage through repeated sequence features. By varying the window size, we can modulate the stringency of this grouping strategy. Small windows produce highly conservative partitions with minimal leakage, while large windows relax this constraint and potentially allow more overlap, enabling robust evaluation of model generalization and sensitivity to local context (Figure 2).

**Figure 2:**
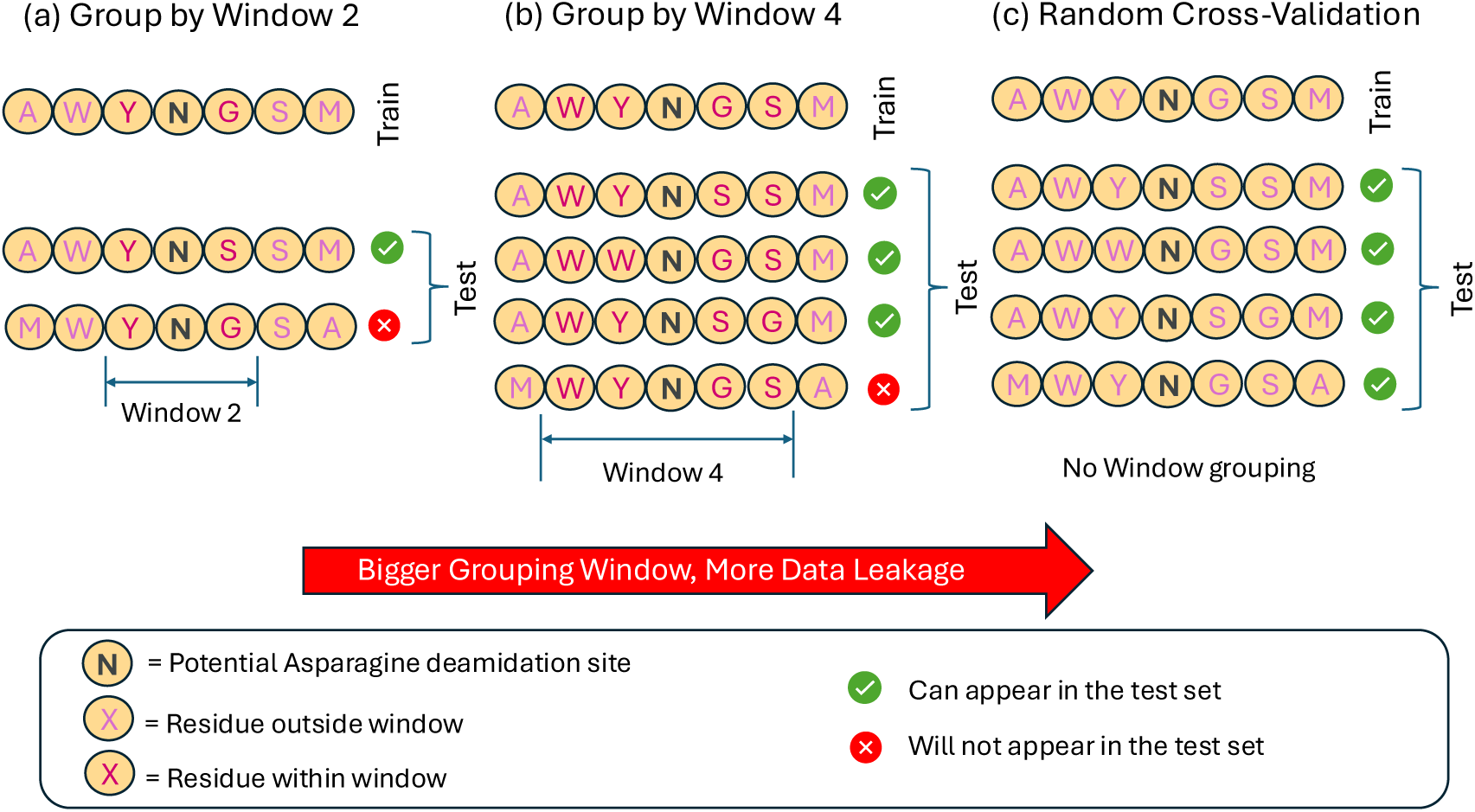
Effect of grouping window on data leakage. In the absence of grouping (C: random cross-validation), overlapping sequence contexts freely occur in both training and test sets, leading to substantial leakage and inflated performance estimates. Grouping reduces leakage by keeping similar windows within the same split; however, as the window size increases, more contexts merge, and residual leakage rises. In the limit of very large windows, grouping converges toward random cross-validation because nearly all contexts overlap.

For each window size and deamidation cutoff, splits are created using StratifiedGroupKFold algorithm [14] which balances both the number of samples and the positive/negative class ratio as much as possible. Additional iterative swaps further refine class balance within a tolerance (see Figure 1b).

Across window sizes 1–6 and both cutoffs (2.0% and 5.0%), the number of unique clusters increases with window size and then saturates (dashed black line in Figure 3; detailed per-cutoff views in Figures 3a and 3b). In all settings, most clusters are negative-only (orange), consistent with the strong class imbalance (few deamidated residues). Mixed clusters (containing both positive and negative samples) decline rapidly as window size grows and reach a stable plateau by window size 4 for both cutoffs. This suggests that a four-residue flanking window is already near-sufficient to separate deamidated from non-deamidated residues; enlarging the window beyond this point adds redundant contextual information without introducing new discriminative patterns (Figure 2). Consequently, the risk of data leakage also plateaus around window size 5.

**Figure 3:**
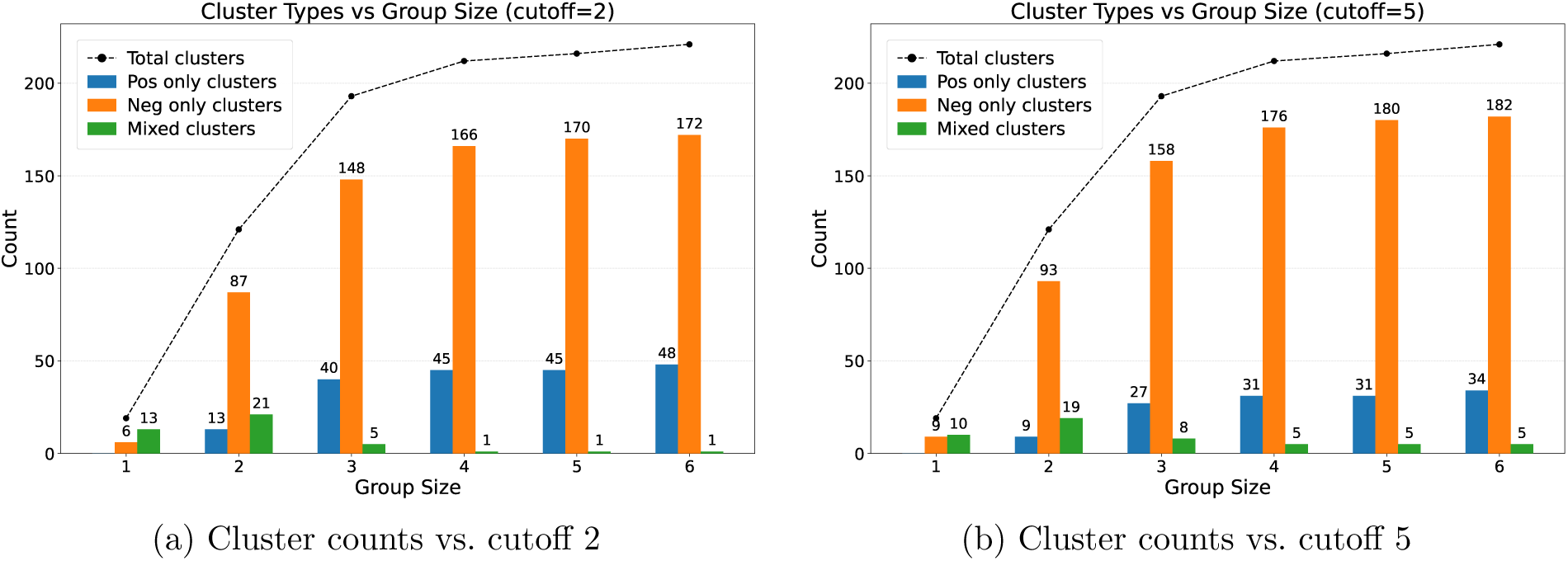
Positive, negative, and mixed cluster counts for grouping sizes 1–6 at cutoffs 2 and 5. Mixed clusters level off by window size 5, indicating a plateau in data leakage. Beyond window 5, no new overlapping patterns appear between train and test.

#### 3.2.2 Nested Cross-Validation

To rigorously assess model performance on data that differ from the training distribution, such as sequences or molecular graphs with novel motifs, domains, or feature combinations not seen during training, we employed nested cross-validation for both the protein language model (PLM) and graph neural network (GNN) pipelines. In each iteration, one fold was held out as the external test set while the remaining data were used for model training and hyperparameter optimization. This approach ensures that evaluation is performed on truly unseen data, providing an unbiased estimate of model generalization.

### 3.3 Overview of Prediction Approaches

#### 3.3.1 Motif-Based Approach

As noted earlier, we use a motif-driven approach grounded in known biochemical patterns of spontaneous asparagine (N) deamidation [18]. In this sequence-level heuristic, a residue is labeled deamidation-prone when an asparagine is immediately followed by glycine (G), serine (S), or asparagine (N). The method is structure-independent, fast, and interpretable, serving as a clear baseline for comparison.

#### 3.3.2 Feature-Based models

A machine learning classifier was used to predict developability properties from selected protein features, as detailed in Section “Protein Structure Generation and Feature Extraction”.

The molecular descriptors included both sequence-based features, such as pphl values and IMGT properties of the N+1 and N–1 residues, and structure-based features, specifically backbone dihedral angles (phi and psi) for the N, N+1, and N–1 residues, as well as asparagine side chain dihedrals (chi1 and chi2), nucleophilic attack distance, solvent accessibility, hydrogen bonding, secondary structure, and local charge. The performance of models using only sequence-based features was compared to those incorporating both sequence- and structure-based features. Notably, sequence-derived features can be computed directly from the protein sequence, obviating the need for structure prediction and substantially reducing computational cost.

To further assess the impact of structural variability and data balance, we experimented with including either the top-ranked AlphaFold2 predicted structure or augmenting the minority class by adding more structural models. Specifically, during training, we balanced the dataset by incorporating the top 25 most probable AlphaFold2-predicted structures for the minority class. Model validation was performed on the same fold split of the validation set without any structural augmentation, allowing for a robust and fair comparison to models relying only on the top-ranked structure.

#### 3.3.3 PLM

We utilize a state-of-the-art protein language model (PLM), ESM-2, which is an evolutionary-scale transformer architecture pretrained on large-scale protein sequence databases [11]. This model captures rich sequence context and biochemical properties across diverse protein families, making it well-suited for protein modeling tasks. To adapt ESM-2 representations to our deamidation prediction task, we explore and systematically compare three distinct training regimes:

- **Frozen PLM:** ESM-2 backbone frozen; train only the downstream MLP classifier.
- **Full fine-tuning:** Train ESM-2 backbone and MLP end-to-end.
- **LoRA fine-tuning:** Parameter-efficient adaptation via low-rank adapters in attention projections, training only small adapter weights while the backbone remains frozen.

In all regimes, contextual residue embeddings are fed into an MLP to predict deamidation probability for target positions, using fixed-size windows around each residue as input.

#### 3.3.4 GNN

To leverage structural context, each protein structure predicted by AlphaFold is converted into a residue-level graph (Figure 4). Nodes correspond to residues (initialized with PLM-derived features); edges encode proximity based on *C_α_* distances within a cutoff *r* and restricted to the *k* nearest neighbors. We evaluate GVP, GAT, and GIN backbones, vary message-passing depth *L*, window size *w*, pooling strategy (mean vs. concatenation), and optimize hyperparameters via nested cross-validation. We report accuracy, precision, recall, F1-score, ROC-AUC, and AUPRC.

**Figure 4:**
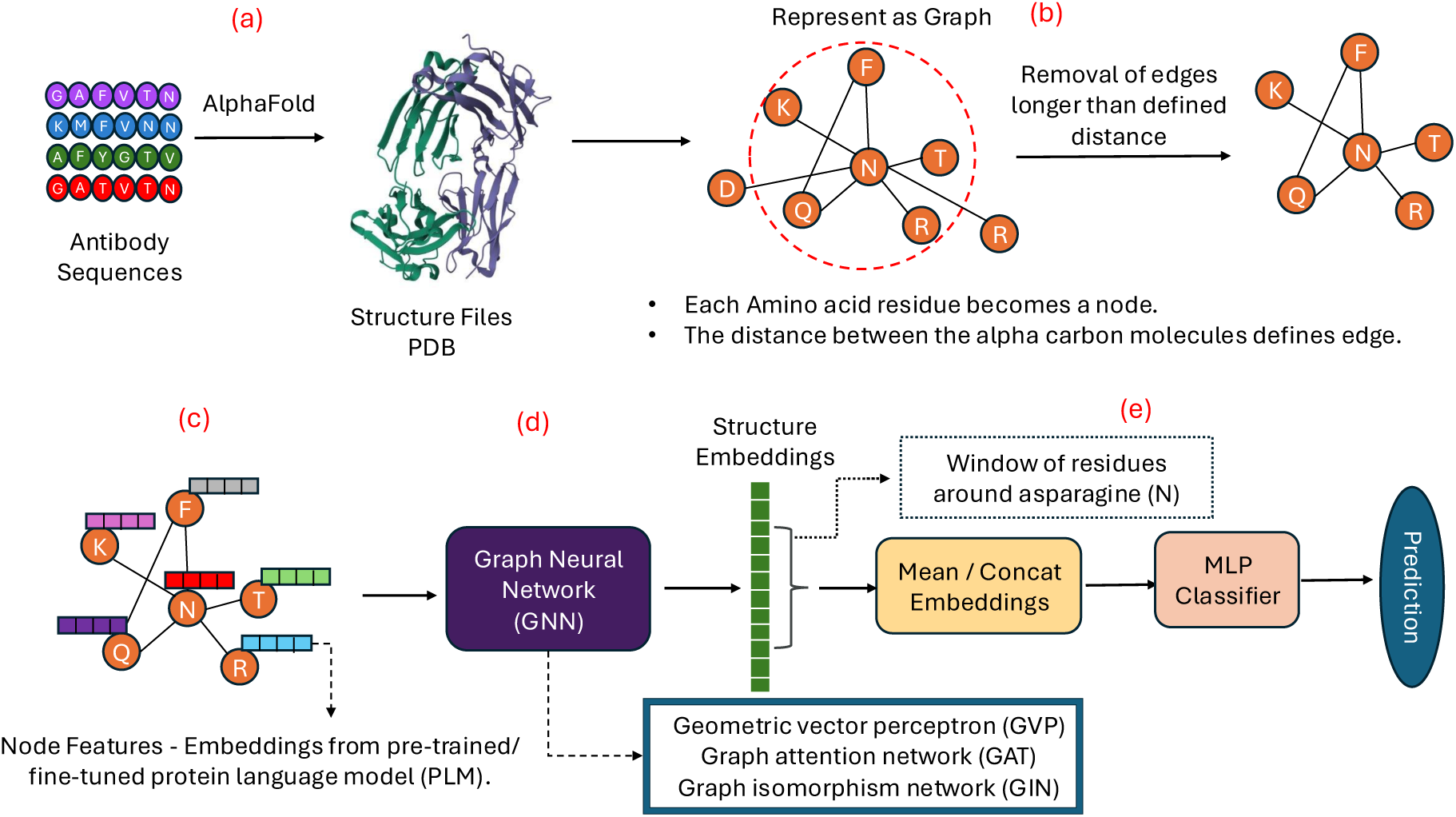
Overview of the GNN architecture for deamidation prediction. (a) Therapeutic protein sequences are first folded via AlphaFold to obtain 3D structural models. (b) The resulting structures are converted into residue-level graphs, where each node represents an amino acid and edges connect residues based on defined spatial distance thresholds. (c) Node features are computed using sequence embeddings generated by a pretrained or a fine-tuned PLM, capturing local biochemical context. (d) These graphs and node features are then processed by a GNN model such as geometric vector perceptron (GVP), graph attention network (GAT), or graph isomorphism network (GIN) to produce structure-aware residue embeddings. (e) For each specified asparagine site targeted for deamidation prediction, a fixed window of neighboring residues is extracted. The embeddings for residues within this window are aggregated and passed through a multi-layer perceptron (MLP) classifier to assess deamidation risk.

**Figure 5:**
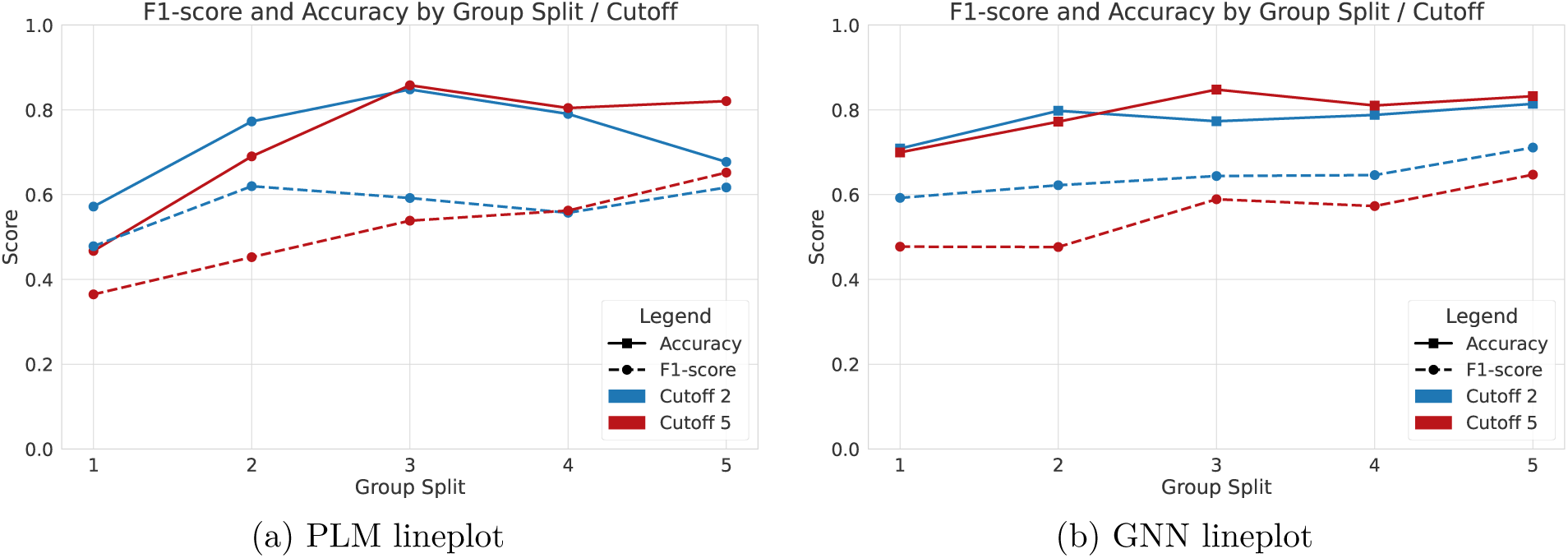
Accuracy and F1-score comparison across group splits (G1–G5) and cutoffs (C2, C5) for both (a) PLM and (b) GNN pipelines.

**Figure 6:**
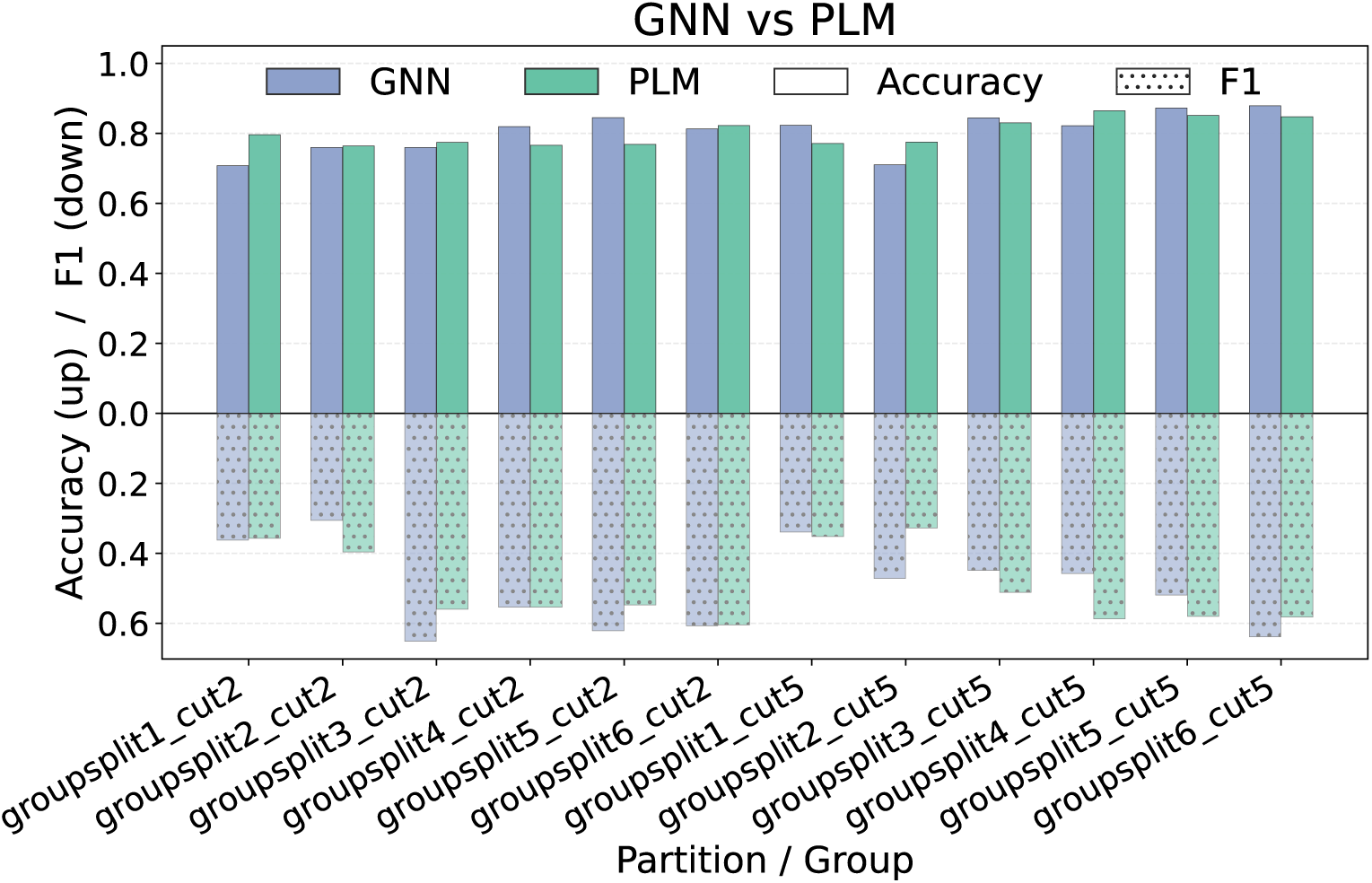
Comparison of the GNN and PLM performance across partitions using nested cross-validation: solid bars (positive axis) show accuracy and hatched bars (plotted downward) show F1-score.

**Figure 7:**
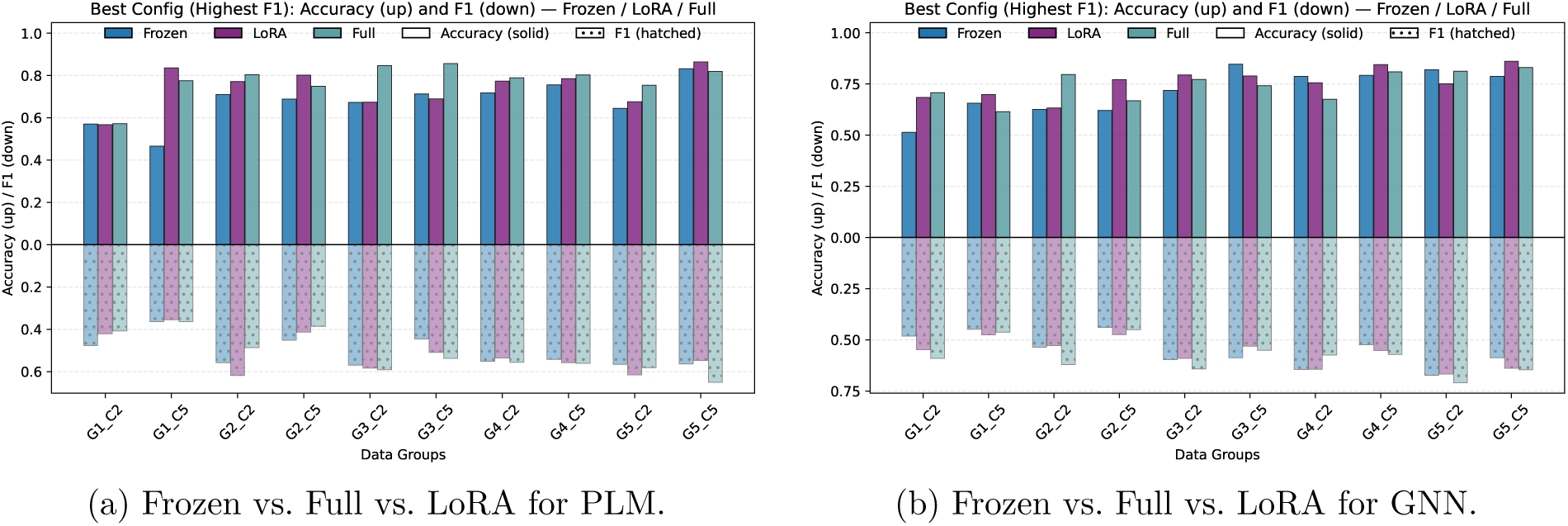
Comparison of frozen, full-parameter, and LoRA fine-tuning for (a) PLM and (b) GNN pipelines based on the highest F1-scores attained per group.

## 4 Results and Comparison

### 4.1 Performance of Motif- and Feature-Based Models

Table 1 reports the performance of rule- and feature-based methods for predicting asparagine deamidation sites. The motif-based baseline achieved moderate accuracy (0.63) and the highest sensitivity (0.63) across models, but its F1-score (0.40) lagged behind feature-driven approaches. A model leveraging sequence-derived features attained higher accuracy (0.76) and AUC (0.66) but exhibited lower F1-score (0.33) and recall (0.37). Introducing structural features markedly improved performance. The Top1 Structure model provided superior accuracy (0.83) and the highest F1-score (0.49). The Top25 Augmented model delivered the highest AUPRC and slightly higher F1-score than Top1 Structure, but with greater variance and higher computational cost.

**Table 1:**
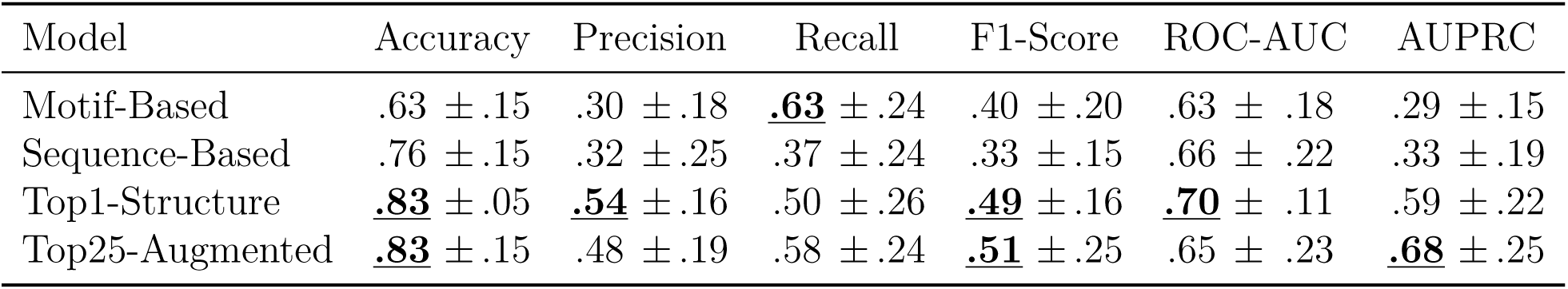
Comparison of cross-validation performance (mean ± standard deviation) for feature-based models on G4_C5. The motif-based baseline relies on sequence patterns, while feature-based models can incorporate both sequence and structural descriptors. Structure augmentation is applied to the minority class. The highest value for each metric is both bolded and underlined.

| Model | Accuracy | Precision | Recall | F1-Score | ROC-AUC | AUPRC |
| --- | --- | --- | --- | --- | --- | --- |
| Motif-Based | .63 ± .15 | .30 ± .18 | **<u>.63</u>** ± .24 | .40 ± .20 | .63 ± .18 | .29 ± .15 |
| Sequence-Based | .76 ± .15 | .32 ± .25 | .37 ± .24 | .33 ± .15 | .66 ± .22 | .33 ± .19 |
| Top1-Structure | **<u>.83</u>** ± .05 | **<u>.54</u>** ± .16 | .50 ± .26 | **<u>.49</u>** ± .16 | **<u>.70</u>** ± .11 | .59 ± .22 |
| Top25-Augmented | **<u>.83</u>** ± .15 | .48 ± .19 | .58 ± .24 | **<u>.51</u>** ± .25 | .65 ± .23 | **<u>.68</u>** ± .25 |

### 4.2 Performance of PLM and GNN Models

Table 2 reports cross-validation performance for PLM and GNN models against Motif-Based and Top1-Structure baselines across group-cutoff-5 splits. Under F1-driven tuning, GNN attains the highest mean F1 in three splits (G2_C5, G4_C5, G5_C5) and is effectively equivalent to the top performer in the remaining two (G1_C5, G3_C5). PLM models often achieve high precision and highest mean accuracy in G3_C5. Top1-Structure achieves single highest mean F1 in G1_C5 and consistently high accuracy but does not consistently outperform GNN on F1. Motif-Based underperforms across nearly all metrics and splits.

**Table 2:** Cross-validation performance (mean ± standard deviation) for GNN, PLM, Motif, and Top1-Structure models across different group–cutoff splits. Gx_Cy denotes the split configuration, where G specifies the grouping scheme and C specifies the deamidation cutoff. For each split, the highest mean value within a metric column is shown in bold and underlined.

| Data | Model | Accuracy | Precision | Recall | F1-Score | ROC-AUC | AUPRC |
| --- | --- | --- | --- | --- | --- | --- | --- |
| G1_C5 | GNN | .70 ± .19 | .38 ± .08 | .76 ± .27 | .48 ± .11 | .69 ± .19 | .39 ± .17 |
| | PLM | .47 ± .13 | .23 ± .08 | **<u>.93</u>** ± .15 | .36 ± .11 | .61 ± .20 | .32 ± .21 |
| | Motif-Based | .65 ± .26 | .17 ± .22 | .57 ± .52 | .25 ± .28 | .63 ± .16 | .25 ± .16 |
| | Top1-Structure | **<u>.80</u>** ± .08 | **<u>.42</u>** ± .20 | .68 ± .31 | **<u>.49</u>** ± .22 | **<u>.76</u>** ± .14 | **<u>.47</u>** ± .19 |
| G2_C5 | GNN | **<u>.77</u>** ± .09 | .39 ± .13 | .66 ± .29 | **<u>.48</u>** ± .19 | **<u>.78</u>** ± .10 | **<u>.48</u>** ± .19 |
| | PLM | .69 ± .24 | **<u>.41</u>** ± .25 | .63 ± .25 | .45 ± .23 | .70 ± .19 | .38 ± .23 |
| | Motif-Based | .63 ± .17 | .27 ± .16 | **<u>.68</u>** ± .38 | .38 ± .22 | .66 ± .18 | .29 ± .08 |
| | Top1-Structure | .76 ± .09 | .36 ± .07 | .40 ± .20 | .35 ± .10 | .62 ± .06 | .35 ± .08 |
| G3_C5 | GNN | .85 ± .05 | .56 ± .17 | .66 ± .25 | .59 ± .17 | **<u>.81</u>** ± .13 | .57 ± .22 |
| | PLM | **<u>.86</u>** ± .06 | **<u>.67</u>** ± .21 | .49 ± .23 | .54 ± .18 | .75 ± .17 | **<u>.58</u>** ± .22 |
| | Motif-Based | .64 ± .09 | .27 ± .13 | .64 ± .30 | .37 ± .17 | .64 ± .13 | .26 ± .11 |
| | Top1-Structure | .79 ± .10 | .49 ± .15 | **<u>.83</u>** ± .09 | **<u>.60</u>** ± .10 | .80 ± .05 | .52 ± .13 |
| G4_C5 | GNN | .81 ± .12 | **<u>.60</u>** ± .23 | .62 ± .18 | **<u>.57</u>** ± .13 | .77 ± .16 | .52 ± .17 |
| | PLM | .80 ± .07 | .50 ± .09 | **<u>.70</u>** ± .23 | .56 ± .11 | **<u>.83</u>** ± .10 | **<u>.62</u>** ± .21 |
| | Motif-Based | .63 ± .15 | .30 ± .18 | .63 ± .24 | .40 ± .20 | .63 ± .18 | .29 ± .15 |
| | Top1-Structure | **<u>.83</u>** ± .05 | .54 ± .16 | .50 ± .26 | .49 ± .16 | .70 ± .11 | .59 ± .22 |
| G5_C5 | GNN | .83 ± .06 | .55 ± .11 | **<u>.85</u>** ± .18 | **<u>.65</u>** ± .09 | **<u>.91</u>** ± .06 | **<u>.74</u>** ± .13 |
| | PLM | .82 ± .18 | **<u>.71</u>** ± .30 | .69 ± .16 | **<u>.65</u>** ± .23 | .85 ± .13 | .69 ± .23 |
| | Motif-Based | .64 ± .11 | .29 ± .17 | .64 ± .31 | .38 ± .20 | .64 ± .18 | .28 ± .11 |
| | Top1-Structure | **<u>.84</u>** ± .05 | .54 ± .11 | .58 ± .21 | .55 ± .15 | .74 ± .11 | .60 ± .23 |

## 5 Discussion

Accurate prediction of asparagine deamidation rates remains a critical challenge in protein therapeutic development. The peptide grouping strategy for cross-validation effectively addresses sequence-similarity-driven data leakage. Our benchmarking across four pipeline architectures highlights the advantages of integrating advanced protein language models (PLMs) and graph neural networks (GNNs), especially for modeling the interplay of sequence and structure that drives degradation susceptibility. Top1-structure delivered strong overall performance among feature-based approaches; PLMs offer scalable, efficient predictions for rapid screening; GNNs integrate explicit three-dimensional information and often deliver the most consistent F1 and ranking gains, particularly under stricter partitions.

## 6 Conclusion

This work presents a systematic approach to predicting asparagine deamidation in protein therapeutics by combining curated datasets, strict evaluation protocols, and deep learning models. Grouping strategies control sequence-similarity leakage, enabling reliable performance assessment. Protein language models provide strong sequence-based predictions, while graph neural networks that incorporate local structural context achieve up to 8% higher accuracy than PLM pipelines and consistent gains over motif-based baselines. Nested cross-validation confirms that these trends are stable across partitioning schemes. The framework is applicable to other degradation pathways and post-translational modifications and supports early-stage in-silico liability assessment.

## Author contributions statement

Conceptualization: N.S., V.S., R.H., J.D., M.P.; Methodology: S.A., N.S., V.S., J.D., M.P.; Data curation: N.S., D.W., J.D.; Model architecture: S.A., M.P.; Validation Strategy Design: S.A., V.S., M.P.; Supervision: J.D., M.P.; Writing—Original Draft: S.A., N.S., J.D., M.P.; Writing—Review and Editing: S.A., N.S., R.J., J.D., M.P.

## Acknowledgments

The authors thank Hang Li, Jun Zhao, Zan Chaudhry, Rupesh Bommana, Auras Bhadra Khanal, Chinmay Gangal, Robert Jones, Jing Ying Yeo, Aarthi Ramsundar, and Yoonji Preville for their support and helpful discussions.

