## Supplement Table 1 for "ACCURATE PREDICTION OF ASPARAGINE DEAMIDATION IN BIOLOGICS USING ADVANCED MACHINE LEARNING MODELS"

**Abstract**

The spontaneous deamidation of asparagine residues remains a major obstacle to the stability and efficacy of protein therapeutics. Currently available models in the literature for predicting deamidation liabilities can suffer from limited generalizability, likely due to biases such as sequence similarity within datasets. In this study, we built machine learning models using pretrained protein language models (e.g., ESM2) and graph neural networks (GNNs), trained on a comprehensive dataset of 591 asparagine sites from over 105 protein molecules. To address the critical issue of data leakage, we implemented a peptide grouping strategy yielding more realistic estimates of model performance for novel deamidation sites. Our analysis shows that, when sequence similarity bias is controlled, protein language models like ESM2 match traditional feature-based models that use amino acid composition, k-mers, PSSMs, and predicted secondary structure/solvent accessibility, while offering substantial computational advantages. Additionally, our GNN-based pipeline further increases prediction accuracy by up to 8% compared to language model-only tools and delivers a 15–25% improvement over motif-based approaches. This methodological framework enables more reliable and rapid in-silico prediction of deamidation liabilities, potentially reducing costly late-stage interventions in protein therapeutic development and is broadly generalizable to the modeling of additional protein post-translational modifications.

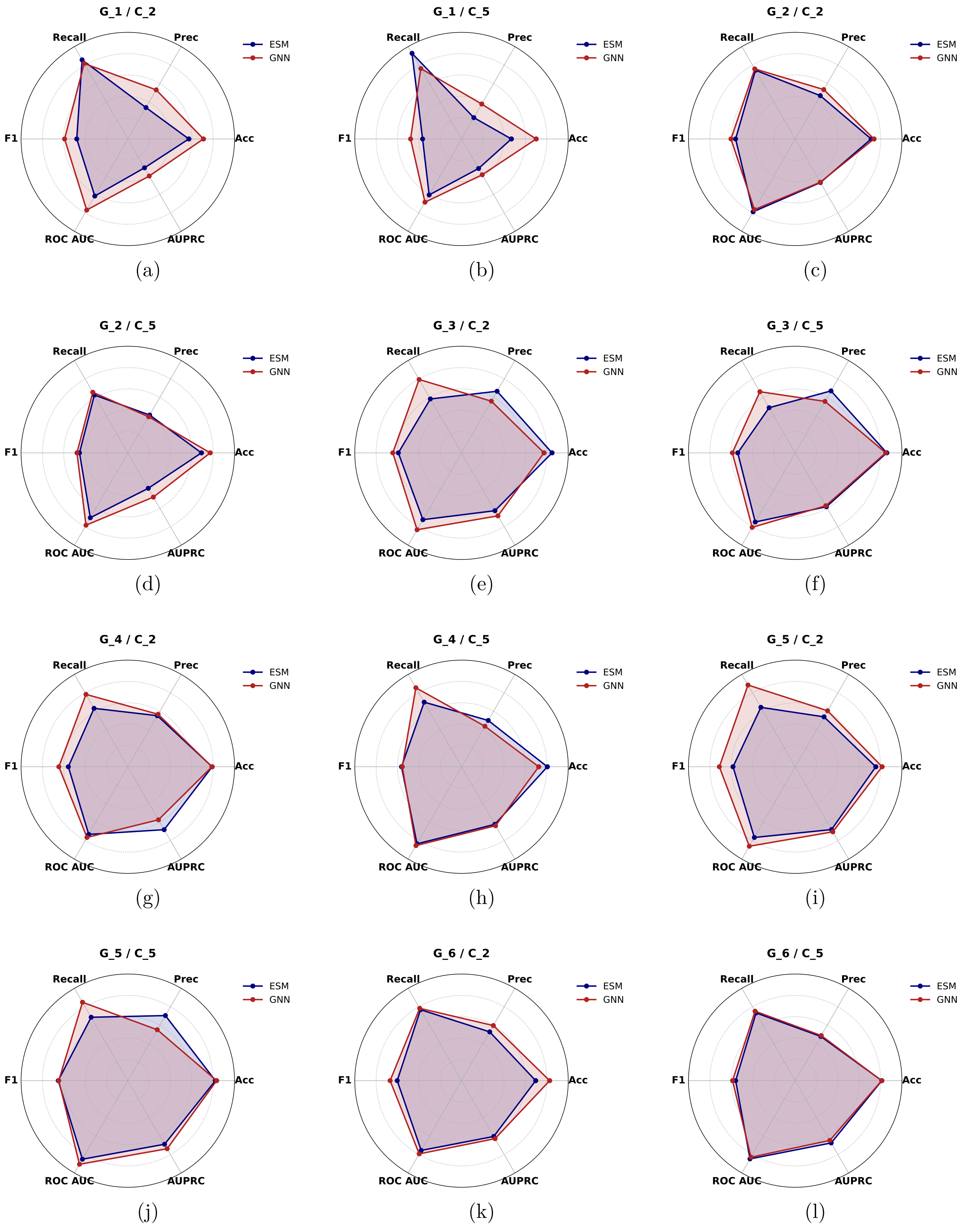


Figure 1: Performance radar plots across grouping windows (G1–G6) and deamidation cutoffs (C2, C5) for multiple metrics.

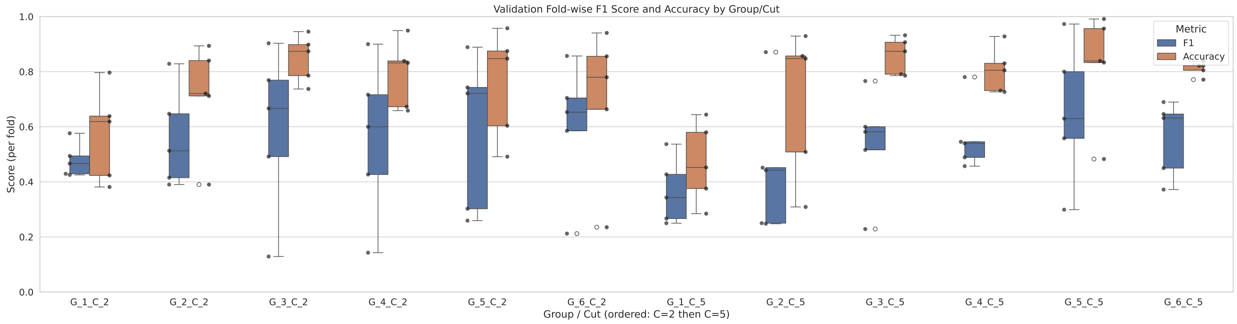


(a) PLM boxplot

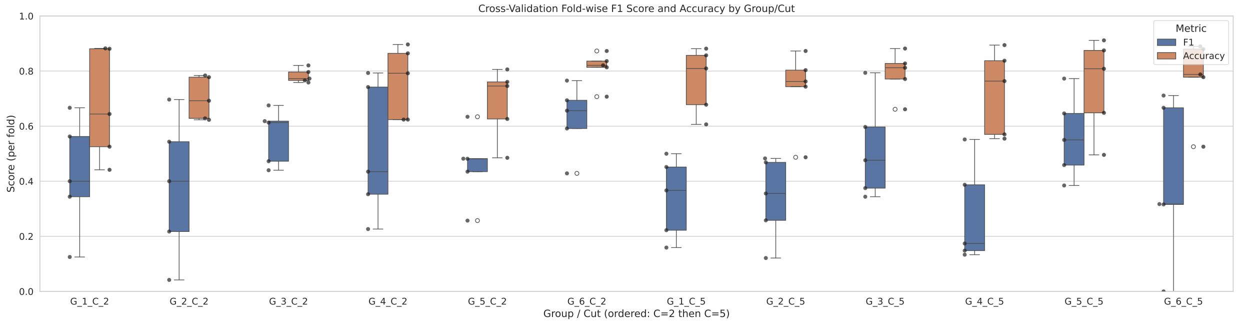


(b) GNN boxplot

Figure 2: Cross-validation F1 and accuracy distributions for PLM and GNN across grouping strategies.

Table 1: Cross-validation performance (mean ± standard deviation) for GNN, PLM, Motif, and Top1-Structure models across different group–cutoff splits. Gx_Cy denotes the split configuration, where G specifies the grouping scheme used to prevent leakage and C specifies the deamidation rate cutoff used to separate good and bad molecules. For each split, the highest mean value within a metric column is shown in bold and underlined for clarity.

| Data | Model | Accuracy | Precision | Recall | F1-Score | ROC-AUC | AUPRC |
|---|---|---|---|---|---|---|---|
| G1_C2 | GNN | **.71** ± .18 | **.53** ± .19 | .82 ± .19 | **.59** ± .09 | **.77** ± .14 | **.77** ± .14 |
| | PLM | .57 ± .15 | .34 ± .07 | **.86** ± .12 | .48 ± .06 | .62 ± .14 | .31 ± .08 |
| | Motif-Based | .68 ± .24 | .25 ± .28 | .56 ± .52 | .33 ± .34 | .64 ± .17 | .33 ± .19 |
| | Top1-Structure | .74 ± .06 | .44 ± .04 | .73 ± .25 | .53 ± .11 | .74 ± .10 | .48 ± .12 |
| G2_C2 | GNN | **.80** ± .09 | **.57** ± .11 | .71 ± .10 | **.62** ± .08 | .75 ± .12 | **.55** ± .19 |
| | PLM | .71 ± .18 | .47 ± .17 | **.74** ± .16 | .56 ± .16 | **.79** ± .10 | .47 ± .15 |
| | Motif-Based | .67 ± .18 | .37 ± .30 | .68 ± .37 | .46 ± .33 | .68 ± .21 | .40 ± .24 |
| | Top1-Structure | .72 ± .07 | .38 ± .23 | .48 ± .30 | .41 ± .26 | .63 ± .15 | .40 ± .17 |
| G3_C2 | GNN | .77 ± .13 | .56 ± .13 | .80 ± .08 | .64 ± .09 | **.83** ± .11 | .68 ± .15 |
| | PLM | **.85** ± .08 | **.67** ± .15 | .58 ± .29 | .59 ± .27 | .72 ± .22 | .63 ± .25 |
| | Motif-Based | .67 ± .08 | .37 ± .15 | .65 ± .27 | .47 ± .19 | .66 ± .13 | .34 ± .14 |
| | Top1-Structure | .81 ± .09 | .61 ± .13 | **.82** ± .11 | **.68** ± .08 | .81 ± .06 | **.76** ± .10 |
| G4_C2 | GNN | .79 ± .13 | **.57** ± .18 | **.78** ± .17 | **.65** ± .16 | **.76** ± .17 | .57 ± .18 |
| | PLM | **.79** ± .11 | .55 ± .23 | .63 ± .30 | .56 ± .26 | .73 ± .24 | **.68** ± .24 |
| | Motif-Based | .67 ± .14 | .40 ± .09 | .69 ± .26 | .48 ± .14 | .67 ± .12 | .35 ± .08 |
| | Top1-Structure | .78 ± .04 | .53 ± .08 | .72 ± .10 | .60 ± .04 | .76 ± .03 | .58 ± .14 |
| G5_C2 | GNN | .81 ± .10 | **.61** ± .17 | **.89** ± .09 | **.71** ± .14 | **.86** ± .08 | **.71** ± .18 |
| | PLM | .76 ± .18 | .54 ± .25 | .64 ± .25 | .58 ± .25 | .77 ± .18 | .68 ± .26 |
| | Motif-Based | .68 ± .06 | .40 ± .07 | .71 ± .26 | .49 ± .11 | .69 ± .09 | .36 ± .07 |
| | Top1-Structure | **.84** ± .05 | .54 ± .11 | .58 ± .21 | .55 ± .15 | .74 ± .11 | .60 ± .23 |
| G1_C5 | GNN | .70 ± .19 | .38 ± .08 | .76 ± .27 | .48 ± .11 | .69 ± .19 | .39 ± .17 |
| | PLM | .47 ± .13 | .23 ± .08 | **.93** ± .15 | .36 ± .11 | .61 ± .20 | .32 ± .21 |
| | Motif-Based | .65 ± .26 | .17 ± .22 | .57 ± .52 | .25 ± .28 | .63 ± .16 | .25 ± .16 |
| | Top1-Structure | **.80** ± .08 | **.42** ± .20 | .68 ± .31 | **.49** ± .22 | **.76** ± .14 | **.47** ± .19 |
| G2_C5 | GNN | **.77** ± .09 | .39 ± .13 | .66 ± .29 | **.48** ± .19 | **.78** ± .10 | **.48** ± .19 |
| | PLM | .69 ± .24 | **.41** ± .25 | .63 ± .25 | .45 ± .23 | .70 ± .19 | .38 ± .23 |
| | Motif-Based | .63 ± .17 | .27 ± .16 | **.68** ± .38 | .38 ± .22 | .66 ± .18 | .29 ± .08 |
| | Top1-Structure | .76 ± .09 | .36 ± .07 | .40 ± .20 | .35 ± .10 | .62 ± .06 | .35 ± .08 |
| G3_C5 | GNN | .85 ± .05 | .56 ± .17 | .66 ± .25 | .59 ± .17 | **.81** ± .13 | .57 ± .22 |
| | PLM | **.86** ± .06 | **.67** ± .21 | .49 ± .23 | .54 ± .18 | .75 ± .17 | **.58** ± .22 |
| | Motif-Based | .64 ± .09 | .27 ± .13 | .64 ± .30 | .37 ± .17 | .64 ± .13 | .26 ± .11 |
| | Top1-Structure | .79 ± .10 | .49 ± .15 | **.83** ± .09 | **.60** ± .10 | .80 ± .05 | .52 ± .13 |
| G4_C5 | GNN | .81 ± .12 | **.60** ± .23 | .62 ± .18 | **.57** ± .13 | .77 ± .16 | .52 ± .17 |
| | PLM | .80 ± .07 | .50 ± .09 | **.70** ± .23 | .56 ± .11 | **.83** ± .10 | **.62** ± .21 |
| | Motif-Based | .63 ± .15 | .30 ± .18 | .63 ± .24 | .40 ± .20 | .63 ± .18 | .29 ± .15 |
| | Top1-Structure | **.83** ± .05 | .54 ± .16 | .50 ± .26 | .49 ± .18 | .70 ± .11 | .59 ± .22 |
| G5_C5 | GNN | .83 ± .06 | .55 ± .11 | **.85** ± .18 | **.65** ± .09 | **.91** ± .06 | **.74** ± .13 |
| | PLM | .82 ± .18 | **.71** ± .30 | .69 ± .16 | **.65** ± .23 | .85 ± .13 | .69 ± .23 |
| | Motif-Based | .64 ± .11 | .29 ± .17 | .64 ± .31 | .38 ± .20 | .64 ± .18 | .28 ± .11 |
